# Investigation of Two-Component Hydrogel System for the Tissue Regeneration from Simulation

**DOI:** 10.1101/2023.12.28.573564

**Authors:** Song Jiang, Ela Kumar

## Abstract

Tissue degenerative diseases pose significant global health challenges. Currently, hydrogel therapy stands as a promising approach to address these conditions. To explore its potential, we developed a computational simulation model to mimic hydrogel behavior accurately and precisely measure its degradation rate while incorporating dynamics associated with cell growth. This model aimed to investigate the relationship between a two-component hydrogel system and cell growth, assessing its feasibility for tissue regeneration. Our analysis revealed that the nutritional support for neural stem cells exceeds that of bone marrow cells, followed by other types of cells. This is significant considering the challenge in culturing neural stem cells compared to the relative ease of culturing other cell types. Additionally, we found that there isn’t a single solution to determine the ‘optimal’ conditions for cell growth. Different tissue regeneration processes require distinct conditions to establish what could be considered as ‘suitable’ growth environments. Lastly, it’s important to note that these ‘suitable’ conditions can be fine-tuned by adjusting various parameters.

## Introduction

Tissue degenerative diseases, such as Parkinson’s disease, dementia, traumatic brain injury, and others, present significant global health challenges.[1] In response, researchers are actively exploring various therapeutic avenues, including drug delivery, cell therapy, gene therapy, and hydrogel therapy, to address these conditions.[2-9] Among these approaches, hydrogel treatment stands out due to its ability to mimic the extracellular matrix, creating a nurturing environment for encapsulated neural stem cells.[10]

Hydrogels, existing in diverse forms and formed through chemical or physical bonding mechanisms, exhibit varying levels of stability. While chemically bonded hydrogels tend to be stable, certain conjugation methods might negatively impact their stiffness and cellular toxicity. Two-component hydrogels has emerged as a promising method due to its rapid formation rate and biocompatible properties among these conjugation methods.[11]

The Two-component hydrogel system is currently recognized, particularly in academia, as one of the most effective biomaterials for tissue regeneration, primarily owing to its outstanding biocompatibility. This system shows the potential to form within milliseconds to seconds by fine-tuning its viscosity within the simulated extracellular matrix environment.[12]

The properties of hydrogels, influenced by numerous factors such as temperature, viscosity, molecular weight, coupling time, and degradation rate, pose challenges for accurate measurement, especially concerning degradation rate determination. The complexity arises due to the interdependence of these various parameters, resulting in an expensive, time-consuming, and labor-intensive measurement process. There are various methods to explore these parameters without resorting to practical experiments, such as statistical analysis, machine learning prediction, and more.[13-16] While these methods offer valuable insights, the development of computational models presents a promising alternative approach in understanding these parameters. In this article, a simulation model has been developed to replicate hydrogel behavior and precisely measure its degradation rate while integrating dynamics related to cell growth.

## Methodology

The dashboard of the system of this model is in the figure 1. It investigates the relationship between two-component hydrogel system with cell growth, and whether it is practicable for tissue regeneration. This system explores the stability of two-component hydrogel with incorporated cells in the micro-biosystem. The ideal micro-biosystem should be that the hydrogel degradation rate is proportional to cell growth rate. The coefficient is determined by many factors, including molecular weight, number of cells, cleavage bonding rate, etc. The micro-biosystem tends to be unstable and is not suitable for biomedical applications if the degradation rate is too slow/fast and cell growth fast is too fast/slow.

**Figure 1.**
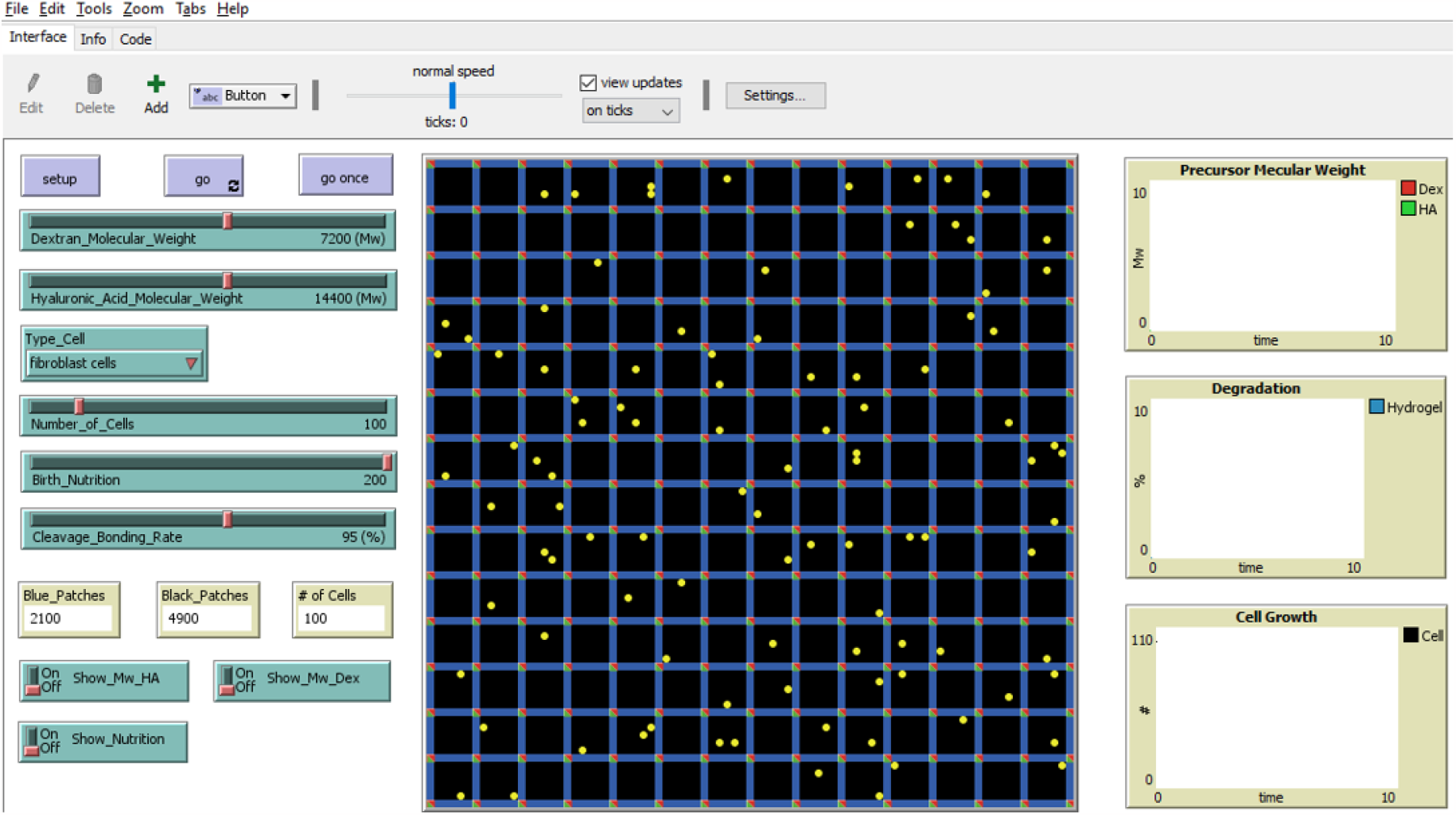
the dashboard of the model

The goal of this model is to provide a brief prediction of hydrogel degradation and cell growth of two-component hydrogel system after a period of time without any labor and cost. Using this model can adjust the hydrogel degradation and cell growth rate to find the most “suitable” conditions for the specific tissue regeneration.

The system entity structure of the model is in the figure 2. The initialized cells will divide after a period of set time. If nutrition is consumed completely, cells will die. If cells bump into unit boundary, they lose some nutrition and rotates in a random degree to continuously move forward. The cells can across the boundary if one or two components are degraded.

**Figure 2.**
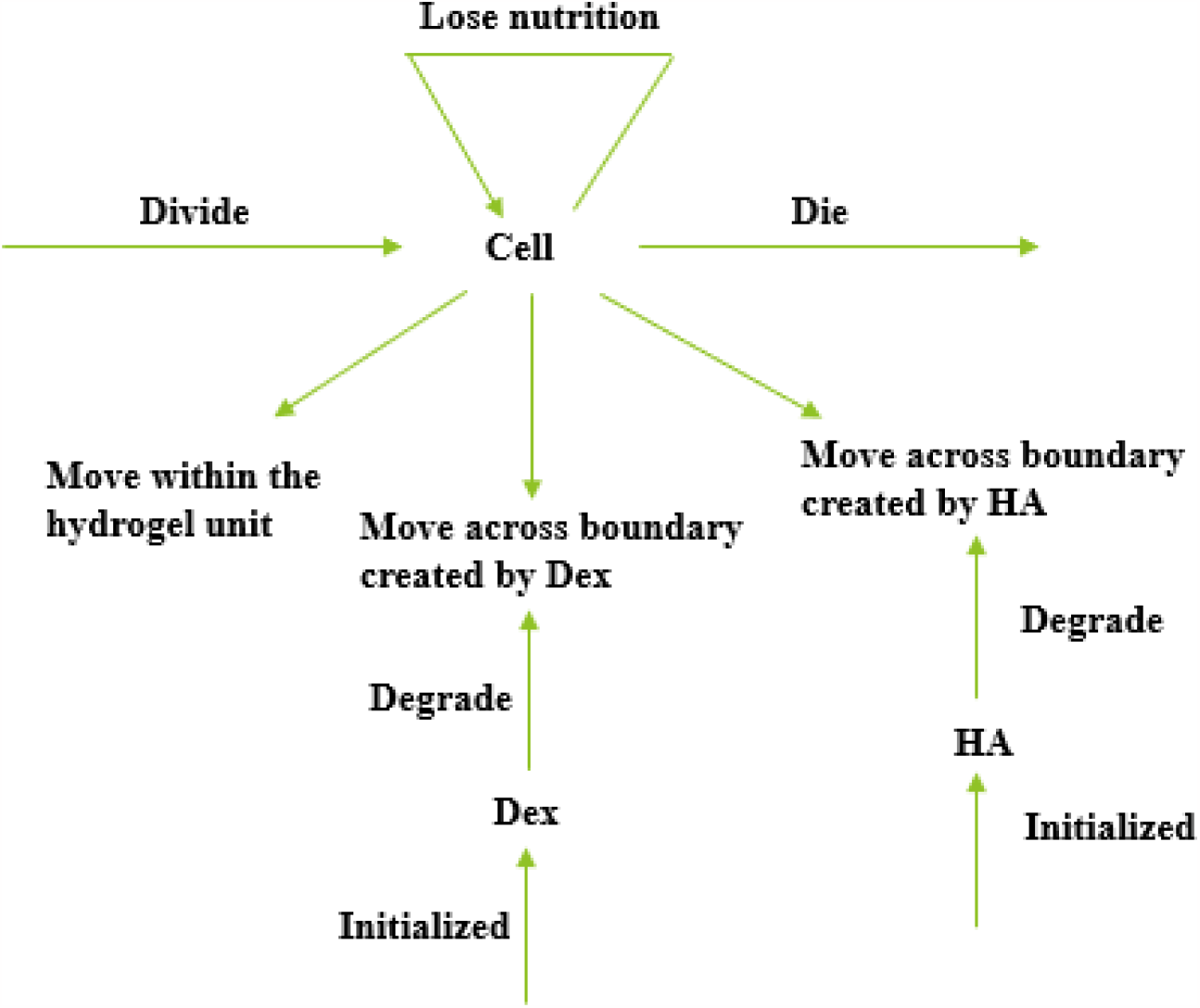
System entity structure of the model

Click the setup button to create the hydrogel networks (green and red) and seed cells (yellow). Click the go button to start this model. It will simulate how this micro-biosystem works. There are five sliders in the controller in the figure 3. Dextran_Molecular_Weight and Hyaluronic_Acid_Molecular_Weight sliders decide molecular weight of two components, respectively. Number_of_Cells slider initiates that how many cells should be seeded. The debonding rate is according to Cleavage_Bonding_Rate slider. The nutrition of cells are firstly given by Birth_Nutrition slider. The type of cells are decided by chooser. When the simulation runs, three plots are exhibited: precursor molecular weight change, degradation rate and cell growth number. Meanwhile, the molecular weight and cell nutrition can be monitored by switching the buttons.

**Figure 3.**
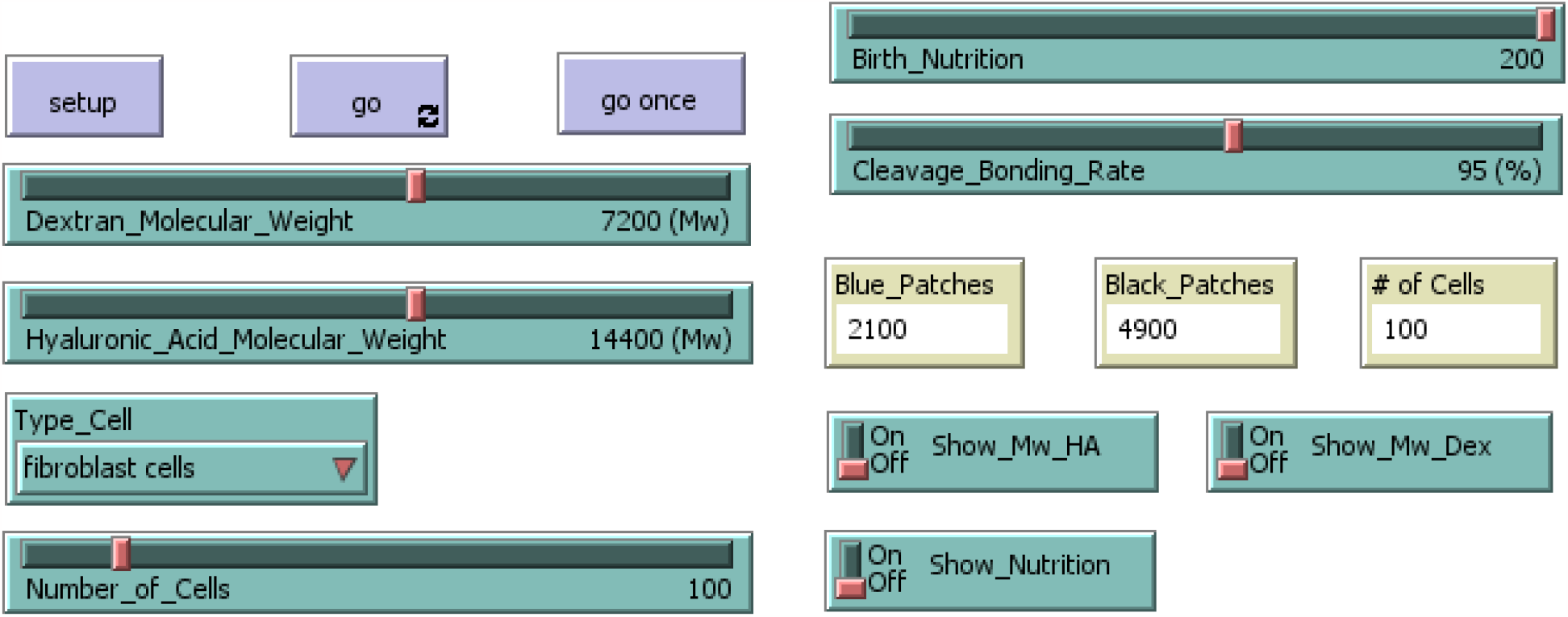
controller of the model

In the beginning of code, three breeds are defined, and values are determined. In this setup function, create-mesh, create-units and Incorporate-cells functions are contained. Create-mesh function is to create hydrogel network. Create-units function is to set hydrogel units. Incorporate-cells function is to seed cells.

In the go, enzyme-degradation, cleavage function, migrate-cells, reproduce-cells, die-cells, die-Dex and die-HA functions are contained. Enzyme-degradation function is to create oriented degradation steps instead of random ones due to the homogeneous hydrogel properties. Cleavage function is to create the degradation made by debonding. Migrate-cells function is to move the cells. Reproduce-cells function is to reproduce the cells. Die-cells function is to wipe out cells with no nutrition. Die-Dex and Die-HA functions are to wipe out hydrogel units totally degraded. Update-globals function is to update degradation rate, Dex and HA molecular weight. In the figure 4, two screenshots are provided. One is initialized result. The second one is the result after running.

**Figure 4.**
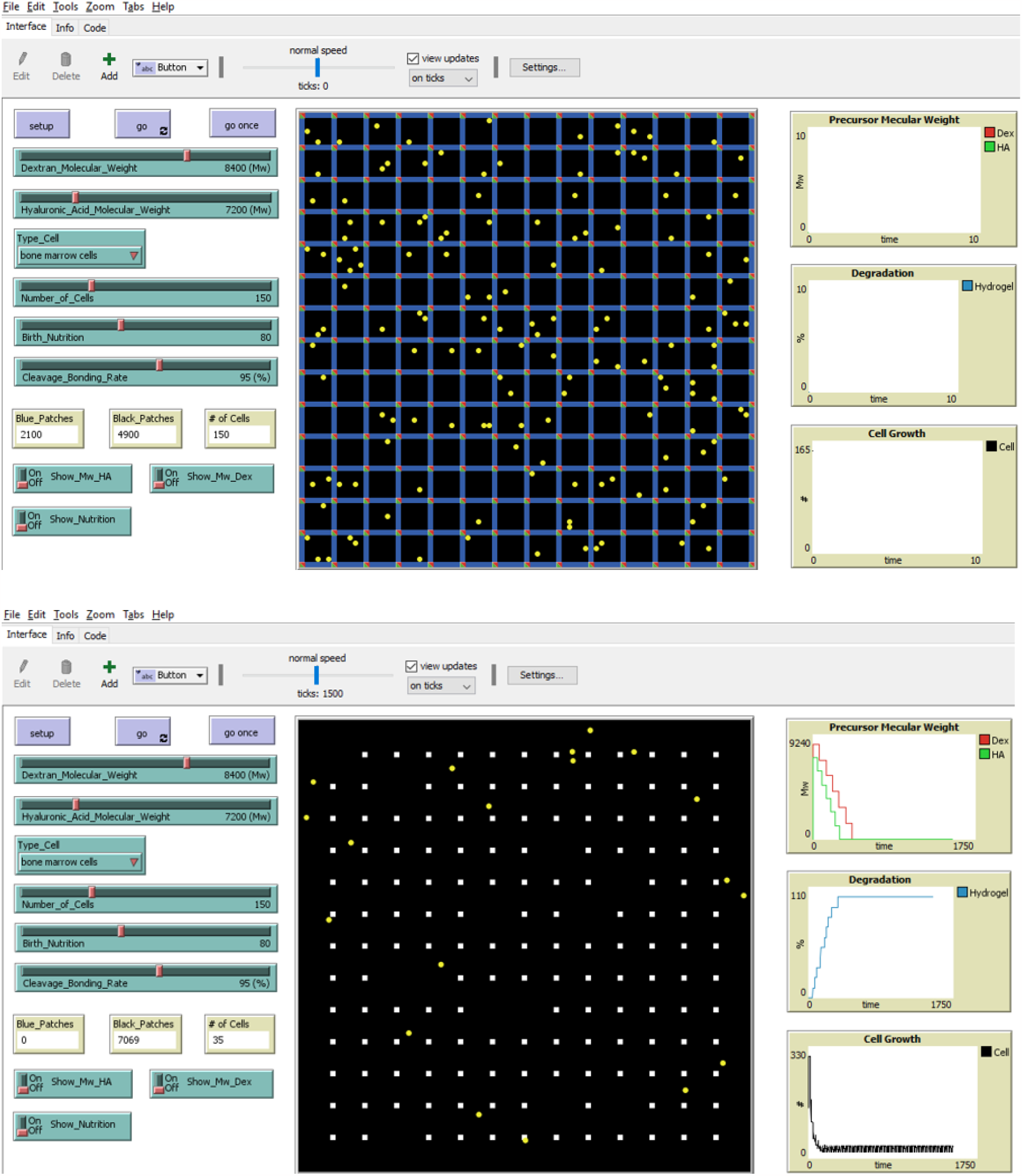
Two snapshots of simulation

**Fig 7:**
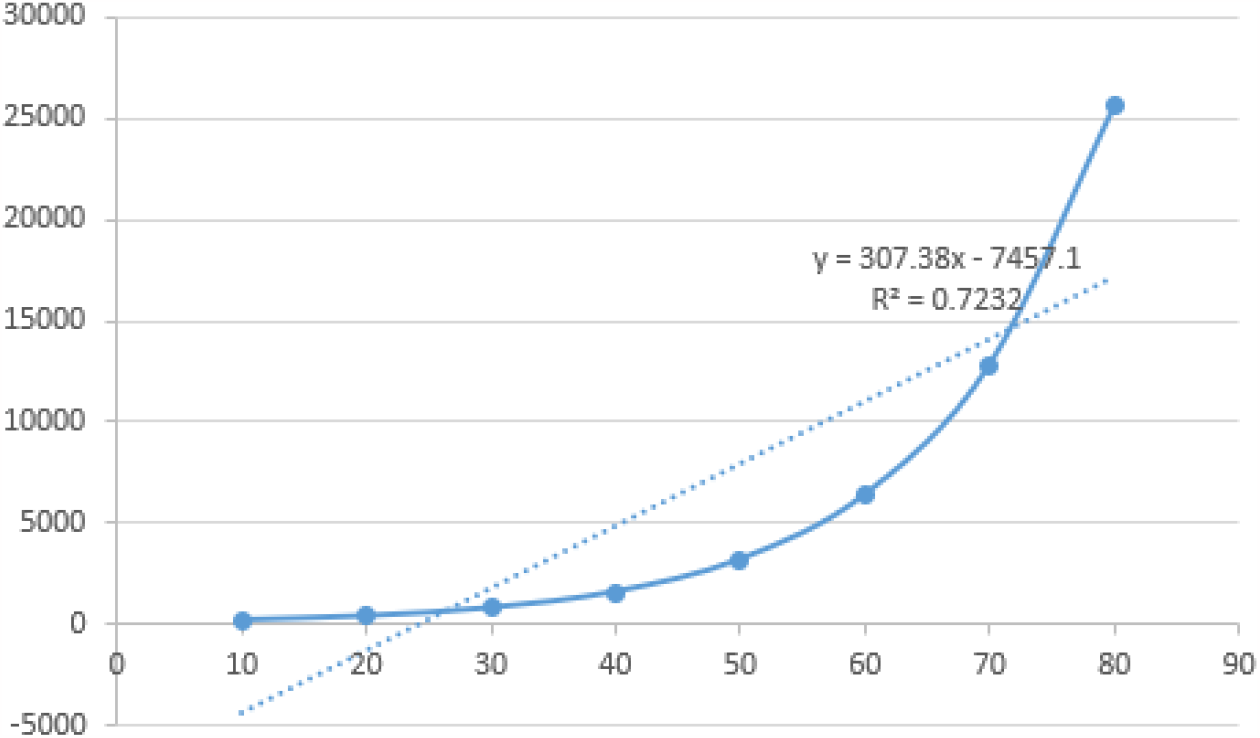
Theoretically, the equation of the number of cells with corresponding hydrogel degradation rate.

**Fig 7:**
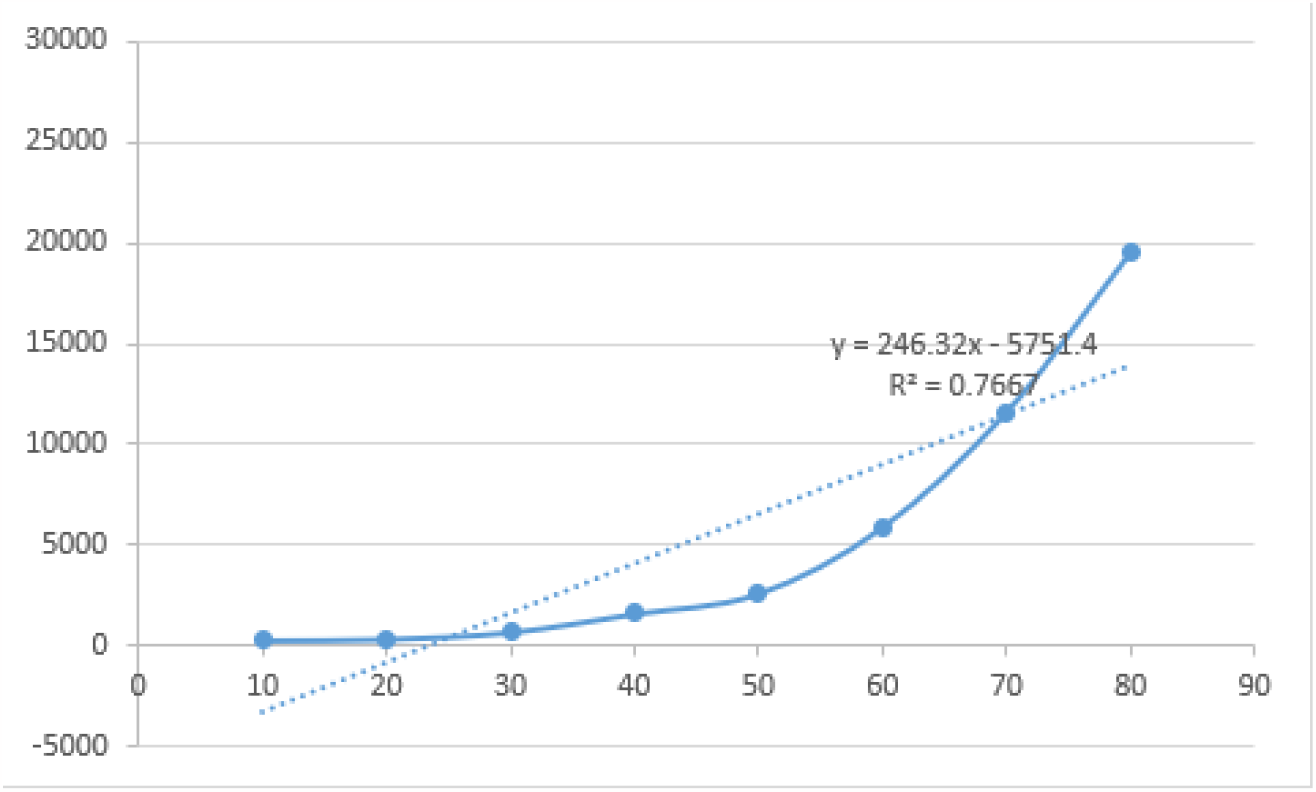
The equation of the number of cells with corresponding hydrogel degradation rate is in the “suitable” conditions for cells.

**Fig 8:**
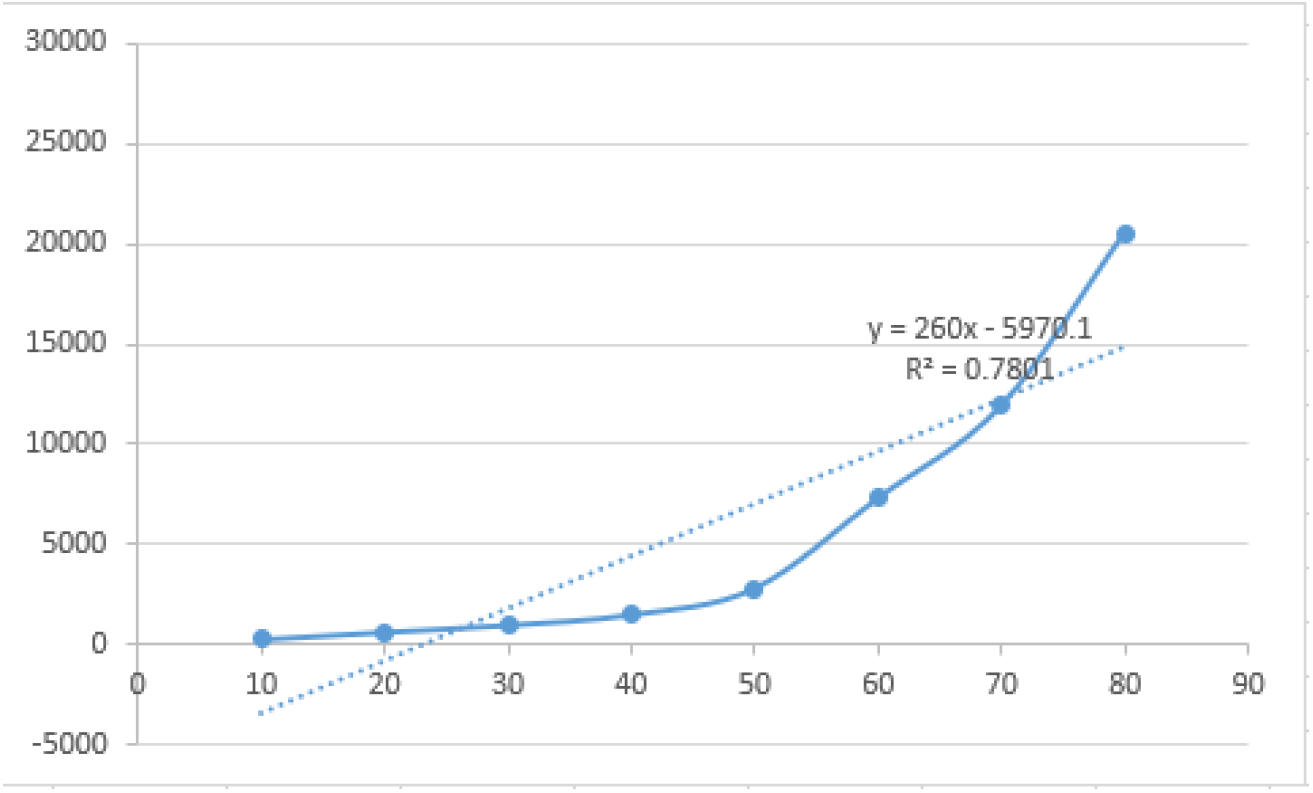
The equation of the number of cells with corresponding hydrogel degradation rate is in the “suitable” conditions for bone marrow cells.

**Fig 9:**
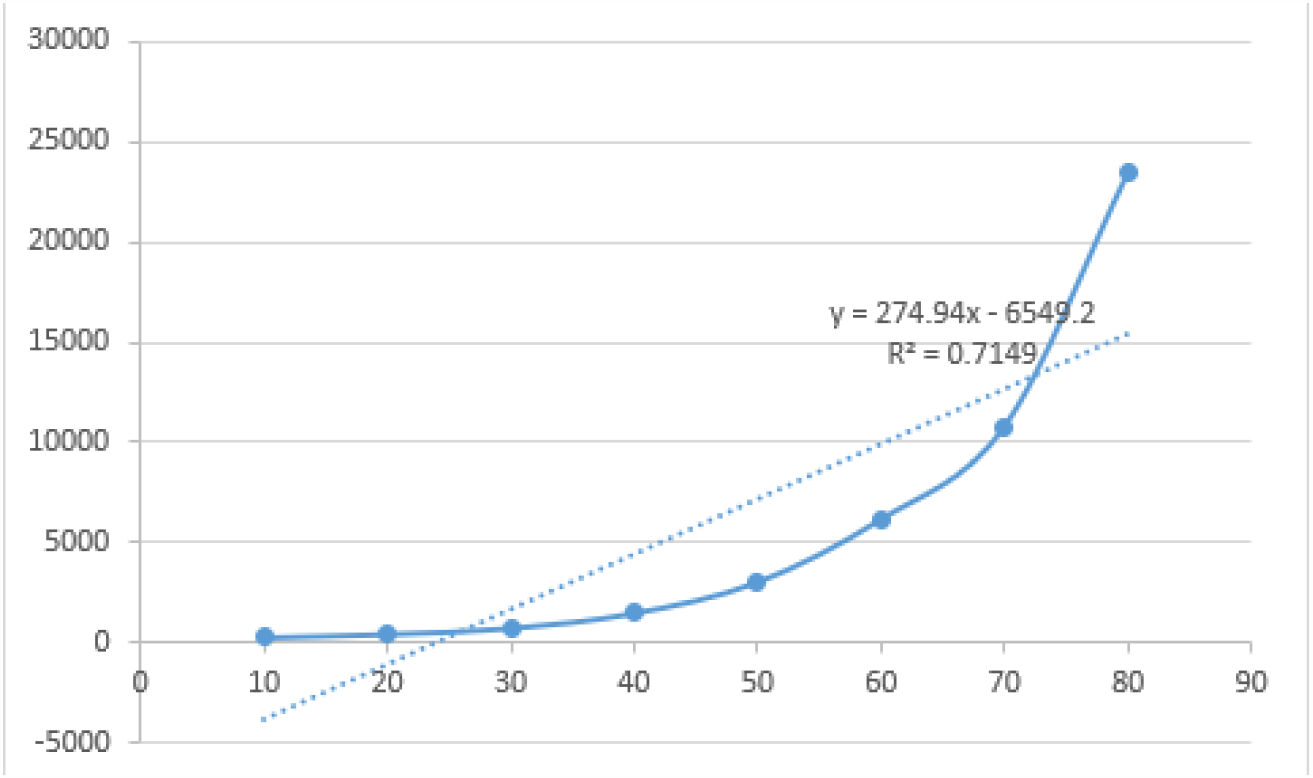
The snapshot of the number of cells with corresponding hydrogel degradation rate is in the “suitable” conditions for neural stem cells.

The model is implemented using 3 breeds of turtle: two for hydrogel units (irremovable), one for cell (movable). The initial number of cells, precursor molecule weight, type cell, etc., are controlled via sliders and chooser. Go once is a one-time procedure. GO is a continuous procedure. When the program is running, these parameters can be adjusted.

### Initialize

- Hydrogel network and units are created.
- Type-cell is decided.
- Number of cells are determined and seeded randomly within the hydrogel network.

### At each tick

- Cell moves 1 step randomly within the hydrogel network units.
- If cell bumps into unit boundary, it loses some nutrition and rotates in a random degree to continuously move forward.
- Cell will divide after a couple of set time.
- Cell will die if the nutrition is consumed totally.
- Cell can across the boundary to move if it degraded.

## Experiments and Results

Before analyzing the results, the “suitable” and “unsuitable” conditions must be defined first. The “suitable” conditions are defined as the cell growth speed is proportional to hydrogel degradation speed. Here, cell specifically divide after every 10% hydrogel degradation. If the initialized number of cells are 100, the degradation rate with corresponding number of cells are in the table below.

**Table 1:** Theoretically, the number of cells with corresponding hydrogel degradation rate is in the “suitable” conditions.

| Degradation rate (%) | 10 | 20 | 30 | 40 | 50 | 60 | 70 | 80 |
| --- | --- | --- | --- | --- | --- | --- | --- | --- |
| # of cells | 200 | 400 | 800 | 1600 | 3200 | 6400 | 12800 | 25600 |

The “unsuitable” conditions are defined as three categories. The first category is that the cell growth speed is much faster than hydrogel degradation speed. Another category is that the cell has no growth when degrading Hydrogel. The last one is that the cell is dead when degrading hydrogel.

The summarized “suitable” and “unsuitable” conditions for three type cells are on the below.

**Table 2:**
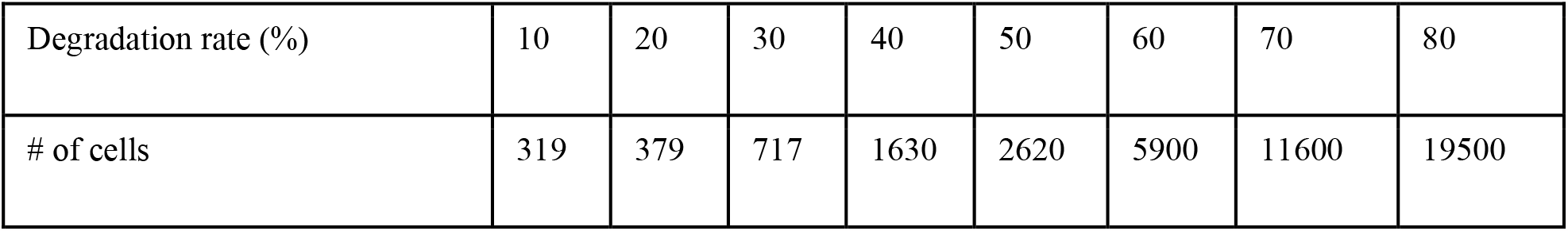
In reality, the number of cells with corresponding hydrogel degradation rate is in the “suitable” conditions for cells.

| Degradation rate (%) | 10 | 20 | 30 | 40 | 50 | 60 | 70 | 80 |
| --- | --- | --- | --- | --- | --- | --- | --- | --- |
| # of cells | 319 | 379 | 717 | 1630 | 2620 | 5900 | 11600 | 19500 |

From above tables, no matter what type of cells, it is obvious that the equations are very similar in the “suitable” conditions. The tested data under the “unsuitable” conditions also match corresponding trends.

**Table 3:** In reality, the number of cells with corresponding hydrogel degradation rate is in the “suitable” conditions for bone marrow cells.

| Degradation rate (%) | 10 | 20 | 30 | 40 | 50 | 60 | 70 | 80 |
| --- | --- | --- | --- | --- | --- | --- | --- | --- |
| # of cells | 256 | 622 | 970 | 1490 | 2800 | 7300 | 11900 | 20500 |

**Table 4:** In reality, the number of cells with corresponding hydrogel degradation rate is in the “suitable” conditions for bone neural stem cells.

| Degradation rate (%) | 10 | 20 | 30 | 40 | 50 | 60 | 70 | 80 |
| --- | --- | --- | --- | --- | --- | --- | --- | --- |
| # of cells | 316 | 488 | 780 | 1530 | 3070 | 6200 | 10800 | 23400 |

**Table 5:** In reality, the cell growth speed is much faster than hydrogel degradation speed in the “unsuitable” conditions for cells.

|  |  |  |  |  |  |  |  |  |
| --- | --- | --- | --- | --- | --- | --- | --- | --- |
| Degradation rate (%) | 10 | 20 | 30 | 40 | 50 | 60 | 70 | 80 |
| # of cells | 810 | 3130 | 6300 | 51600 | too big to run | too big to run | too big to run | too big to run |

**Table 6:** In reality, the cell has no growth when degrading hydrogel in the “unsuitable” conditions for cells.

|  |  |  |  |  |  |  |  |  |
| --- | --- | --- | --- | --- | --- | --- | --- | --- |
| Degradation rate (%) | 10 | 20 | 30 | 40 | 50 | 60 | 70 | 80 |
| # of cells | 200 | 156 | 141 | 126 | 126 | 126 | 126 | 126 |

**Table 7:** In reality, all cells are dead in the “unsuitable” conditions for bone marrow cells.

|  |  |  |  |  |  |  |  |  |
| --- | --- | --- | --- | --- | --- | --- | --- | --- |
| Degradation rate (%) | 10 | 20 | 30 | 40 | 50 | 60 | 70 | 80 |
| # of cells | 0 | 0 | 0 | 0 | 0 | 0 | 0 | 0 |

## Conclusion

From analysis, it is obvious that neural stem cell nutrition given is larger than bone marrow cell, followed by cell. Because neural stem cell is hardest cell to culture, whereas cell is easiest one. Moreover, the solution to find the “suitable” conditions for cell growth is not unique. Also, different tissue regeneration needs different requirements to create the “suitable” conditions. In the last, “Suitable” conditions can be adjusted by tuning the paraments.

## Future Work

In reality, the nutrition given is from medium. That is not set by different type cells need different medium. To make the model more realistic, the parameter of nutrition should be added in the model to make this system simulate degradation and cell growth more precisely and accurate.

